# Stumpy forms are the predominant transmissible forms of *Trypanosoma brucei*

**DOI:** 10.1101/2023.08.30.555461

**Authors:** Jean Marc Tsagmo Ngoune, Parul Sharma, Aline Crouzols, Nathalie Petiot, Brice Rotureau

**Affiliations:** Trypanosome Transmission Group, Trypanosome Cell Biology Unit, Institut Pasteur, Université Paris Cité, INSERM U1347, Paris, France; Sorbonne Université, ED515 Complexité du Vivant, Paris, France; Parasitology Unit, Institut Pasteur of Guinea, Conakry, Guinea

**Keywords:** *Trypanosoma brucei*, stumpy, slender, tsetse, transmission

## Abstract

Schuster *et al*. demonstrated that bloodstream slender forms of African trypanosomes are readily transmissible to young tsetse flies where they can complete their complex life cycle (1). In their experimental conditions, a single slender parasite was sufficient for productive infection. Here, we compared the infectivity of slender and stumpy bloodstream forms in adult flies with a mature immune system, and without using any chemical compounds that would alter the insect immune response and/or promote the infection. After ingestion of slender forms, infected flies were observed only in one out of 24 batches of non-immunocompetent teneral flies and with a high number of parasites. In contrast, infected flies were detected in 75% (18/24) of the batches infected with stumpy parasites, and as few as 10 stumpy parasites produced mature infections in immune adult flies. We discuss that, although Schuster *et al*. have demonstrated the intrinsic capacity of slender form trypanosomes to infect young and naive tsetse flies, highlighting the remarkable plasticity and adaptability of these protists, this phenomenon is unlikely to significantly contribute to the epidemiology of African trypanosomiases. According to both experimental and field observations, stumpy forms appear to be the most adapted forms for African trypanosome transmission from the mammalian host to the tsetse fly vector in natural conditions.

## Introduction

Protist parasites of the *Trypanosoma brucei* group cause Human African Trypanosomiasis (HAT), or sleeping sickness in humans, and nagana in cattle (2). They are transmitted by the blood feeding tsetse fly following a long (at least 3 weeks) and complex (at least 9 distinct stages) cyclical development (review in (3)). In the mammalian host’s blood circulation, proliferating slender trypanosomes differentiate into cell cycle-arrested stumpy cells upon quorum sensing when they reach high parasite densities (4–7). This differentiation is thought not only to regulate the parasite load in the reservoir host (8), but also to provide transmissible parasites adapted to pursue the life cycle in the vector host (9). Indeed, stumpy forms express several transcripts and proteins necessary to the next developmental stage in the insect, the procyclic form, including the Protein Associated to Differentiation 1 or PAD1 (5). For decades, arrest of the cell cycle and differentiation to the stumpy stage were presumed essential for the developmental progression of bloodstream trypanosomes to the insect stages.

Recently, Schuster *et al*. demonstrated that slender trypanosomes can also present some intrinsic characteristic of transmissible forms (PAD1 mRNAs and proteins) and are readily transmissible to both young male and female tsetse flies, where they can complete their complex life cycle (1), yet with a lower efficiency than stumpy forms (10). In their experimental conditions, a single slender parasite was sufficient for productive infection. These laboratory conditions are however significantly different from what is encountered in the field. First, only young teneral flies (1-3 days post-eclosion) with an immature immune system were used. Second, in some experiments, chemical compounds altering the insect immune response (glutathione) and/or promoting the infection (N-acetyl-glucosamine) were added to the infective meal. To assess the importance of these parameters, we challenged the infectivity of slender bloodstream forms in adult tsetse flies, i.e. in conditions closer to the natural situation.

## Material and methods

### Strains, culture and *in vitro* differentiation

The AnTat 1.1E Paris pleomorphic strain of *Trypanosoma brucei brucei* was derived from a strain originally isolated from a bushbuck in Uganda in 1966 (11). Bloodstream form trypanosomes were cultivated in HMI-9 medium supplemented with 10% (v/v) FBS (12) at 37°C in 5% CO_2_. Proliferative slender cells were maintained at densities lower than 5.10^5^ parasites/ml to prevent their natural quorum-sensing-dependent differentiation into stumpy forms. For *in vitro* slender to stumpy BSF differentiation, we used 8-pCPT-2′-O-Me-5′-AMP, a nucleotide analogue of 5’-AMP (BIOLOG Life Science Institute, Germany). Briefly, 2×10^6^ pleomorphic AnTat 1.1E slender forms were incubated with 8-pCPT-2′-O-Me-5′-AMP (5 μM) for 48 h (13). Freshly differentiated stumpy forms and slender cells were then centrifuged at 1,400 × g for 10 minutes and resuspended at the appropriate densities in SDM-79 medium supplemented with 10% FBS. Cells were resuspended at either 10^3^, 10^4^ or 10^5^ parasites / ml. Assuming individual bloodmeal volumes ranging between 10-100 μl, this would correspond to ingestions of 10-100, 100-1,000 or 1,000-10,000 parasites per condition.

### Tsetse fly maintenance, infection and dissection

*Glossina morsitans morsitans* tsetse flies were maintained in Roubaud cages at 27°C and 70% hygrometry and fed through a silicone membrane with fresh mechanically defibrinated sheep blood (BCL, France). Adult (between 2 and 3 weeks after emergence) or teneral males (between 24h and 72h post-emergence) were allowed to ingest parasites through a silicone membrane. No chemical supplement was used in the first set of experiments. For assessing the effect of immunomodulatory compounds in the second set of experiments, 60mM *N*-acetylglucosamine were added to the infective meal. A total of 3 to 5 independent biological replicates per condition were performed with batches of 50 flies per condition.

Flies were starved for at least 24 hours before being dissected blindly 28 to 31 days post-ingestion for isolation of all stages from the midgut and salivary glands. For recovery of all tsetse organs, after rapid isolation of the salivary glands in a first drop of phosphate buffer saline (PBS), whole tsetse alimentary tracts, from the distal part of the foregut to the Malpighian tubules, were dissected and arranged lengthways in another drop of PBS as previously described (14, 15). Isolated organs were then scrutinized under a microscope at 40x magnification by two independent readers and infection rates per organ were scored (16).

### Immuno-fluorescence analysis (IFA)

Cultured parasites were washed in TDB and spread onto poly-L-lysine coated slides. For flash methanol fixation, slides were air-dried for 10 min, fixed in methanol at −20°C for 5 seconds and rehydrated for 20 min in PBS. For immunodetection of stumpy forms, slides were incubated for 1 h at 37°C with a rabbit polyclonal anti-PAD1 antibody (kindly provided by Keith Matthews, University of Edinburgh) (5) diluted at 1:300 in PBS containing 0.1% Bovine Serum Albumin (BSA). After 3 consecutive 5 min washes in PBS, a species and subclass-specific secondary antibody coupled to the Alexa 488 fluorochrome (Jackson ImmunoResearch) diluted at 1:1000 in PBS containing 0.1% BSA was applied for 1 h at 37°C. After washing in PBS, slides were finally stained with 4’,6-diamidino-2-phenylindole (DAPI, 1 µg/ml) for visualization of kinetoplast and nuclear DNA content and mounted under coverslips with ProLong antifade reagent (Invitrogen), as previously described (14). Slides were observed under an epifluorescence DMI4000 microscope (Leica) with a 100x oil objective (NA 1.4) to assess the proportion of PAD1-positive cells in the infective meals (n > 100 cells / condition).

## Statistical analysis

Infections rates were compared by a two-sided ANOVA at 95% confidence with Prism V10.0.3 (GraphPad). MG infection rate comparisons were statistically significant between teneral and adult flies infected with ST in each amount (p<0.02 with 10 parasites; p<0.0001 with 100 and 1,000 parasites) and with 1,000 SL (p<0.0001). MG infection rate comparisons were statistically significant (p<0.0001) between parasite stages (SL and ST) in each amount (10, 100 and 1,000) and for each fly group (teneral and adult), excepted in teneral flies infected with 1,000 parasites (p=0.2356).

## Results

Pleomorphic *T. b. brucei* bloodstream forms were either maintained in culture at a density lower than 5.10^5^ parasites / ml to prevent quorum-sensing-induced differentiation and obtain only slender forms or induced for differentiation with a 5’- AMP nucleotide analogue to obtain mostly stumpy forms. The expression of PAD1 at the cell surface was assessed by immunofluorescence analysis prior to each experimental infection: no PAD1 expression was detected in the slender group, whereas an average of 63% (52% to 71%, n = 12 replicates) of the induced cells were expressing PAD1. Batches of 50 teneral (< 72h) or adult (2-3 weeks) male tsetse flies were fed in parallel with either slender or stumpy forms at densities corresponding to individual ingestion of about 10, 100 or 1,000 parasites per bloodmeal. In total, 1,384 flies from 12 distinct experimental infections were dissected about 4 weeks (28 to 31 days) after parasite ingestion. Infection rates in midguts and salivary glands were quantified and plotted for each condition (Figure 1 and Table S1).

**Figure 1.**
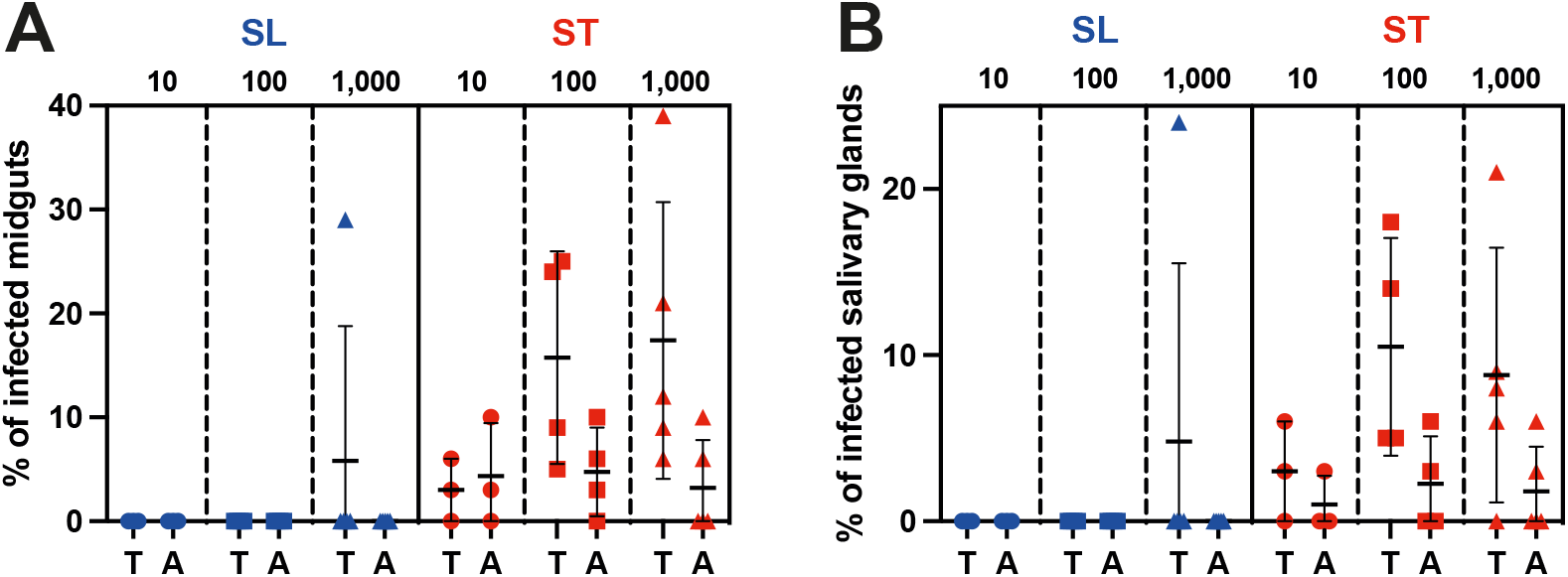
Stumpy forms are the predominant transmissible forms. Comparison of (**A**) midgut and (**B**) salivary gland infection rates in teneral (T) Vs. adult (A) tsetse flies (batches of 50 flies) infected with 10 to 100 (circles, 3 independent experiments), 100 to 1,000 (squares, 4 independent experiments) or 1,000 to 10,000 (triangles, 5 independent experiments) parasites in the slender (SL in blue) or stumpy (ST in red) forms.

After ingestion of slender forms, infected flies were observed in only 1 batch out of 24. This occurred in not yet fully immunocompetent teneral flies and with the highest number of ingested parasites (1,000 to 10,000 parasites). In contrast, midgut and salivary glands infected flies were observed in 75% (18/24) and 62.5% (15/24) of the batches infected with stumpy parasites, respectively. As few as 10 stumpy parasites produced mature infections in immunocompetent adult flies and the infection rates were similar whatever the amounts of stumpy forms ingested. In more susceptible non-immune teneral flies, the infection rates were increasing with the number of stumpy forms ingested.

Differences between the strain clones, the cell culture conditions and/or the fly colony maintenance conditions could explain part of the differences in infection rates observed here as compared to the Schuster *et al*. study (1). Nevertheless, the use of the lectin- inhibitory sugar N-acetyl-glucosamine to enhance infection rates in the latter study could be a more likely explanation. To assess this hypothesis, an additional experimental challenge was performed to compare infection rates in teneral versus adult flies, with or without N-acetyl-glucosamine supplement in an infective meal containing 10^5^ slender parasites / ml (Figure 2). Whereas no infection was detected in adult flies, the N-acetyl-glucosamine supplementation of the infective meal led to an increase of the infection rates from 2,4% to 13,3% in teneral flies (Figure 2).

**Figure 2.**
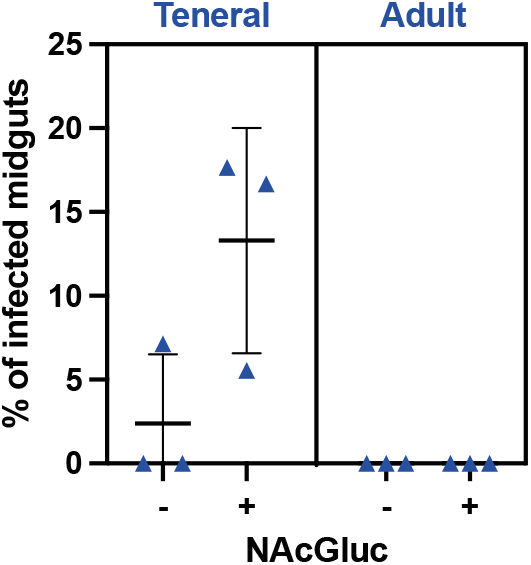
*N*-acetylglucosamine promotes trypanosome infection in teneral flies. Comparison of midgut infection rates in teneral Vs. adult tsetse flies (batches of 50 flies) infected with 1,000 to 10,000 slender parasites with (+) or without (−) N-acetyl- glucosamine (NAcGluc) supplement in the infective meal containing 10^5^ slender parasites / ml (equivalent to 1,000 to 10,000 slender parasites per meal).

## Discussion

The findings of Schuster *et al*. (1) have opened a debate on the traditional view of the trypanosome life cycle where slender trypanosomes are considered as non-competent for cyclical development in the insect vector (18). The authors proposed that their observations could provide a solution to a long-lasting paradox, namely the successful transmission of parasites in chronic infections, despite low parasitemia. Nonetheless, Schuster *et al*. performed all their experimental infections in laboratory conditions that were optimized for maximum transmission efficiency, with the use of teneral flies, and for some experiments, the addition of *N*-acetylglucosamine or glutathione (1).

Teneral flies are young flies that remain unfed for up to 3-5 days after emergence from their puparium. They are known to be significantly more susceptible to trypanosome infection as multiple studies demonstrated they are immunologically immature (weak immune system and, leaky peritrophic matrix) (19–22). According to capture-recapture studies (23), teneral flies however represent a minority of individuals in wild tsetse populations. Hence, knowing that adults can live up to 9 months (24), the impact of teneral flies on trypanosome transmission may be limited, if not incidental.

In addition, Schuster *et al*. supplemented most infective meals with 60 mM *N*- acetylglucosamine, an inhibitor of tsetse midgut lectins (17) that was also confirmed to enhance trypanosome infectivity in teneral flies in the present study. For infections with monomorphic parasites, the addition of 12.5 mM glutathione, an antioxidant that reduces the midgut environment, protected trypanosomes from cell death induced by reactive oxygen species (25). In total, these two chemical compounds used in Schuster *et al*. (1) have inhibited the immune response in teneral flies and substantially enhanced the chances for slender trypanosomes to develop in the insect vector.

Conditions are far less favourable in the field; hence we investigated and compared the infection potential of slender and stumpy forms in adult and teneral flies without the addition of chemicals. We observed that slender forms were not infective to adult tsetse flies and only at densities higher than 10^5^ parasites / ml. Nevertheless, in endemic areas, especially in Western Africa, parasitemia in confirmed cases are usually very low (< 10^4^ parasites / ml in Guinea for instance) and it is necessary to concentrate parasites in blood prior to microscopic examination to increase sensitivity of parasitological diagnosis (2). Hence, the possible transmission of a few slender trypanosomes from the blood of individuals with a chronic infection is unlikely to explain the maintenance of the parasite circulation in tsetse populations.

By contrast, we observed that as few as 10 stumpy parasites are enough to produce mature infections in both teneral and adult flies, already with a significant efficiency. In patients with low parasitemia, the quorum-sensing-triggered differentiation of slender to stumpy forms could be compatible with, or even more adapted to, extravascular forms present in some tissues and organs, with a limited dilution of the parasites and parasite factors remaining concentrated locally. Indeed, extravascular PAD1-positive trypanosomes were detected in high numbers at least in adipose tissues (26) and in the dermis (27) of experimentally infected mice. The presence of PAD1-positive extravascular trypanosomes was also assessed in the skin of confirmed gambiense HAT cases and unconfirmed seropositive individuals in endemic areas (28) (and unpublished data). This suggests that stumpy trypanosomes accessible to tsetse flies are likely more abundant than previously estimated in individuals with low parasitemia.

Schuster *et al*. have demonstrated the intrinsic capacity of slender form trypanosomes to infect young and naive tsetse flies, highlighting the remarkable plasticity and adaptability of these protists. The fine understanding of the underlying cellular mechanisms and / or transient adaptations involved in this process remains an exciting challenge. This event is however unlikely to contribute to the epidemiology of African trypanosomiases in natural settings. According to both experimental and field observations, stumpy forms appear to be the most adapted forms for African trypanosome transmission from the mammalian host to the tsetse fly vector in natural conditions.

## Acknowledgements

We thank K. Matthews for providing the anti-PAD1 antibody and P. Bastin for his critical reading of the manuscript.

## Author contributions

AC, NP, JMTN and PS performed the experiments. BR designed the study, analysed the data, and wrote the manuscript. JMTN, PS and BR discussed the manuscript.

